# Identification of Kinases Activated by Multiple Pro-Angiogenic Growth Factors

**DOI:** 10.1101/2021.02.28.433132

**Authors:** Scott Gruver, Scott Rata, Leonid Peshkin, Marc W Kirschner

## Abstract

Antiangiogenic therapy began as an effort to inhibit VEGF signaling, which was thought to be the sole factor driving tumor angiogenesis. It has become clear that there are more pro-angiogenic growth factors that can substitute for VEGF during tumor vascularization. This has led to the development of multi-kinase inhibitors which simultaneously target multiple growth factor receptors. These inhibitors perform better than monotherapies yet to date no multi-kinase inhibitor targets all receptors known to be involved in pro-angiogenic signaling and resistance inevitably occurs. Given the large number of pro-angiogenic growth factors identified, it may be impossible to simultaneously target all pro-angiogenic growth factor receptors. Here we search for kinase targets, some which may be intracellularly localized, that are critical in endothelial cell proliferation irrespective of the growth factor used. We develop a quantitative endothelial cell proliferation assay and combine it with “kinome regression” or KIR, a recently developed method capable of identifying kinases that influence a quantitative phenotype. We report the kinases implicated by KIR and provide orthogonal evidence of their importance in endothelial cell proliferation. Our approach may point to a new strategy to develop a more complete anti-angiogenic blockade.

## Introduction

Nearly all tissues require vascularization to maintain homeostasis. Angiogenesis, or the formation of new microvasculature from existing vessels, allows for tissue growth in both normal and pathological circumstances (Potente, Gerhardt, and Carmeliet 2011). A few examples include much of development, wound healing, the formation of the placenta during pregnancy, diabetic retinopathy, and cancer. The complex process involves the coordination of many growth factor-dependent processes including matrix degradation, cell proliferation, motility, morphogenesis, and apoptosis. It was originally thought that vascular endothelial growth factor, or VEGF, was necessary and sufficient for both normal and pathological angiogenesis (Carmeliet 2005). Indeed, blocking VEGF signaling can greatly improve clinical outcomes in the pathological cases of diabetic retinopathy (Crawford et al. 2009) and macular degeneration (Cabral et al. 2017). However, given the somewhat disappointing failure of therapies targeting VEGF in cancer, it appears that, at least in the case of tumor angiogenesis, other growth factors can suffice in the absence of VEGF signaling (Jászai and Schmidt 2019; Khan and Bicknell 2016; Grépin and Pagès 2010).

Renal cell carcinoma (RCC), the most common form of kidney cancer, is one particular cancer where the inhibition of angiogenic signaling is under heavy investigation (Hsieh et al. 2017). RCC provides an instructive example of the importance of VEGF and other angiogenic growth factors. RCC patients treated with VEGFR2 kinase domain inhibitors show moderate increased survival but also eventually develop resistance (Grünwald and Merseburger 2013; Choueiri et al. 2015). Upregulation of other pro-angiogenic growth factors, namely HGF and FGF, have been suggested as potential factors leading to such resistance (Mollica et al. 2019; Zhou et al. 2016). Interestingly, newer generation VEGFR inhibitors which also possess the ability to inhibit other pro-angiogenic factors provide clinical benefit even in patients with total resistance to VEGFR2 mono-inhibition (Choueiri et al. 2015). This has led to calls for the development of a more complete angiogenic blockade (Grünwald and Merseburger 2013). Given the lack of direct comparative studies on the pathways used by different pro-angiogenic growth factors, it is currently unclear what a complete angiogenic blockade would require.

Growth factors have many effects on cells, with perhaps none so well appreciated as their requirement for the passage of untransformed cells through the cell cycle (Gross and Rotwein 2016). Upon the completion of mitosis, if cells in culture find themselves in the absence of growth factors, they exit the cell cycle and enter into a reversible quiescent state (Zetterberg and Larsson 1985). If growth factors are subsequently replaced, cells re-enter the cell cycle, resume growth, duplicate their genome, and divide. The intracellular signaling pathways that transduce the growth factor signal to cell cycle machinery have been intensively studied and many key signaling pathways have been identified for specific growth factors. However, many cell types, such as endothelial cells, respond to multiple growth factors, raising the question of uniqueness and redundancy in growth factor signaling pathways.

In the absence of growth factors, untransformed cells with intact apoptotic pathways also become apoptotic (Sarkar and Mandal 2009; Letai 2006). Thus, the addition of growth factors also serves to suppress pro-apoptotic signaling and cell death. From this perspective, growth factors are often referred to as survival factors (Collins et al. 1994). The mechanisms by which individual angiogenic factors promote passage through the cell cycle and suppress death have been studied to a certain extent. How growth factor signaling pathways comparatively influence cell proliferation is less well studied.

We present here a quantitative assay to study the intracellular signaling responses to proangiogenic growth factors with the goal of identification of shared pathways in primary dermal human microvascular endothelial cells (DMECs). We focus on how growth factors influence proliferation, or the growth of endothelial cell populations. Proliferation is a complex phenotype determined by a balance of birth and death. The assay requires the formulation of a basal proliferation medium in which a low rate of proliferation is achieved through the balance of a low birth rate and a low, but non-zero, death rate. The assay is used to screen ~30 reported angiogenic growth factors for effects on proliferation and identify three that produce a robust increase in proliferation. We then assay the effects of a carefully chosen panel of kinase inhibitors, which when combined with KIR, a recently developed machine learning method (Taranjit Singh Gujral, Peshkin, and Kirschner 2014; Rata et al. 2020; Taranjit S. Gujral et al. 2014), implicates specific kinases as important in DMEC proliferation in response to specific growth factors. Focusing on the intersection of the implicated kinases in each growth factor tested, we identify kinases that are central to endothelial cell proliferation regardless of the growth factor input. Finally, we apply an orthogonal method to contribute to the evidence for the kinases implicated by KIR.

### Establishment of a Quantitative Endothelial Cell Proliferation Assay

We wish to create an assay and accompanying analytical framework that will produce quantitative measures of DMEC proliferative response to single purified pro-angiogenic growth factors. This includes generating an experimental system that allows for the passive counting of cells over time, computing proliferation rates and characterizing the proliferative response of DMCEs to pro-angiogenic growth factors. We then use the assay to determine the quantitative effects of a highly characterized set of kinase inhibitors on DMECs’ proliferation rate. Inhibitor alteration of the proliferation rate is then fed into KIR analysis, generating a ranked list of kinases with predicted contribution to setting the proliferation rate.

#### The Experimental System

To create a quantitative, time-lapse assay of proliferation in response to purified growth factor, we labelled the nuclei of primary dermal microvascular endothelial cells with nuclear-localized mCherry (Fig1A). Imaging of NLS-mCherry-labeled DMECs counterstained with a DNA-binding dye revealed the labelling efficiency to be ~97% (not shown). Using 384-well plates, with a single 4x image, we were able to image the entire population of labeled cells within a well. Expression of the nuclear-localized mCherry offered the ability to use image processing methods to count cells in real-time (FigS1A). Through comparison with manually counted images, we estimated an absolute counting error of 3% (not shown). The counting error from the automated method was independent of cell density (Fig.S1B). Finally, expression of this nuclear reporter proved to be passive as it did not alter the doubling time when compared to unlabeled parental population (Fig S1C). Therefore, we are able to count cells proliferating *in vitro* in a reliable, non-invasive manner.

**Figure 1.**
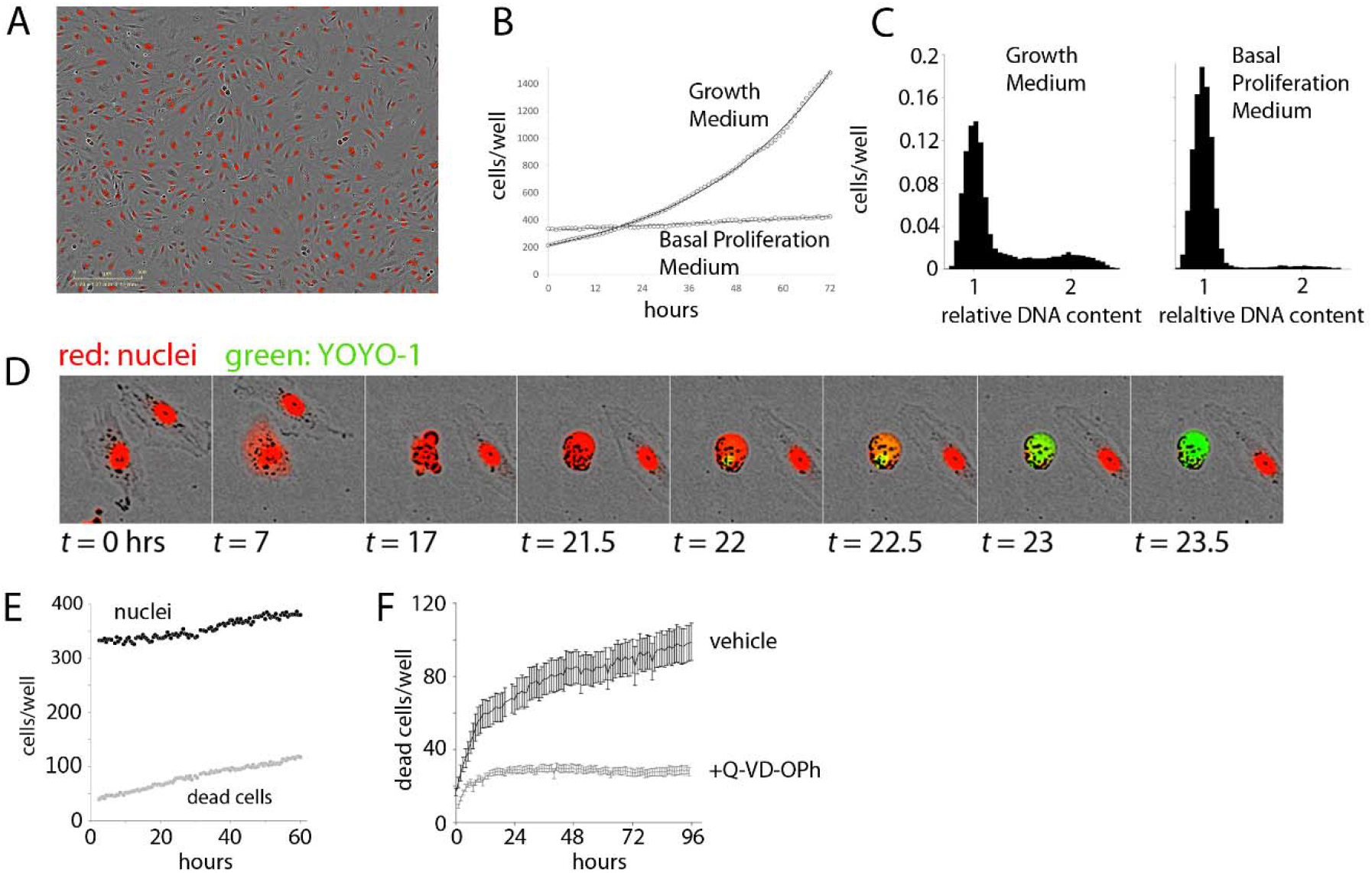
Establishment of the Experimental System. **A.** Demonstration of Dermal Microvascular Endothelial Cell (DMEC) labeling. Fluorescent image of nuclear-localized mCherry overlaid on a phase contrast image of confluent DMECs. B. The proliferation of DMECs in full growth medium (dotted line) is exponential (solid line is a fitted exponential curve). **C.** Cell cycle analysis of relative DNA content determined by labeling live cells with Hoescht and imaged with a microscope. Full growth medium (left) produces a familiar distribution of cells while 24 hours in Basal Proliferation Medium (BPM) (right) produces far fewer cells in S- and G2/M-phases. D. Demonstration of YOYO-1 dye to detect dead cells. E.) Example of nuclei (closed circles) and dead cells (open circles) over time in BPM. F. Cell death in BPM is countered by inhibition of caspase activity via 10 μM Q-VD-OPh.

#### The Proliferative Behavior of Cells in the Experimental System

Next, we examined the proliferative behavior of DMECs after plating in full growth medium. Full growth medium contains 5% FBS and a cocktail of growth factors. Following an approximately 24-hour period of attachment in this rich growth medium (Fig. S1D), the cells entered a phase of exponential growth (Fig 1B). Proliferation continued exponentially for another 70 hours, deviating only as the confluency reached ~70%. Such exponential growth is expected in a population where there are abundant resources (excess growth factors), and inhibitory crowding effects (contact inhibition) are absent (see Box1). These results also defined the confluency below which contact inhibition remains insignificant and we were able to directly verify that in the presence of growth factors, the proliferation rate is independent of plating density (Fig, S1E).

###### Box 1

Cell proliferation, or growth of cell population, combines the rates of birth and death. The *per capita* proliferation rate, P, is the difference between the birth rate (*B*) and death rate (*D*)

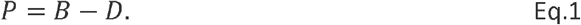

It is worth specifically pointing out some implications of the fact that the proliferation rate is determined by a balance of cell birth and death. On the extreme end of the spectrum where conditions are entirely favorable, the death rate will be zero and *P* = *B*. Conversely, when conditions do not allow for birth but also lead to death, *P* = -*D*. In conditions in between these two extremes, any observed proliferation rate can be achieved through a balance of birth and death.

The per capita birth rate has units of births per cell per unit time while the death rate has units of deaths per cell per unit time. As our experimental system allows for the measurement of live and dead cells over time, we wish to obtain estimates of birth and death rates through time-lapse measurements of the number of nuclei and the number of dead cells. To do so, we first define the proliferation rate as:

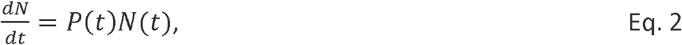

where *N*(*t*) is the number of nuclei at time *t* and *P*(*t*) is the instantaneous proliferation rate. The generality of this simple model is apparent since from it one can arrive at a whole family of commonly applied models of population growth. For example, if *P*(*t*) is defined as a constant, say *k_P_* then equation 2 has the solution of exponential growth:

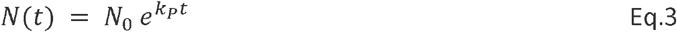

Where *N*_0_ is the population size at time *t* = 0. Equation 3 well describes many populations, particularly in the early phase of growth (i.e., before density-dependent effects begin to dominate). Likewise, if one assumes a density dependence of the population growth of a specific form, then equation can be solved to yield the logistic growth model.

Since equation 2 depends on *P*(*t*), the issue therefore becomes a matter of discovering what *P*(*t*) looks like in our assay. Equation 2 can be rearranged to

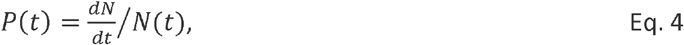

to obtain a useful form for examination of P(t) from the data as everything on the right-hand side is easily observable. Similarly, we can study the death rate by defining it as

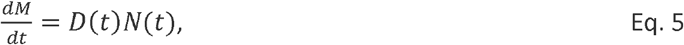

where *M*(*t*) is the number of dead cells at time *t* and *D*(*t*) is the time-dependent death rate. Equation 5 can be rewritten in the same manner as equation 2 to reveal a useful formula for the death rate.

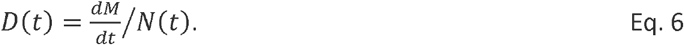

Armed with *P*(*t*) and *D*(*t*), we can then easily obtain *B*(*t*) from Equation 1, i.e.,

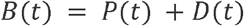

Having characterized the proliferative response in the presence of excess growth factors, we next attempted to isolate the proliferative effects of purified growth factors. To accomplish this, we formulated a basal proliferation medium (BPM) without any exogenous growth factors and significantly reduced serum. The goal of this medium was three-fold: 1.) to increase the dynamic range of the assay by a reduction in the basal proliferation rate compared to that observed in full growth medium, 2.) to achieve specificity through a reduction in background growth factor signaling due to their presence in serum, and 3.) to model the re-entry of resting endothelial cells in vessels into the cell cycle by synchronization of cells in the G0 phase of the cell cycle. Ideally, cells in a basal medium would show little to no net proliferation. We explored the possibility of achieving this goal via titration of serum in a commonly used endothelial basal medium (Zetterberg and Larsson 1985). When the serum content was too low (i.e., < 0.5%) a net loss of cells occurred, suggesting that many cells died from growth factor withdrawal. From these experiments we defined basal proliferation medium (BPM) to contain 1% serum. Comparing the proliferative response of DMECs in BPM (Fig. 1B, E), where there is very little net increase in the number of cells over time, to full growth medium (Fig. 1B), it is clear BPM offers considerable dynamic range.

Resting endothelial cells in a blood vessel, before receiving pro-angiogenic stimuli, would be in a quiescent state, or the so-called G0 phase of the cell cycle. Using DNA-staining for cell cycle analysis and comparing DMECs in full growth medium and in BPM shows the effects of BPM on the cell cycle (Fig. 1C). Specifically, in growth medium, the majority of cells are found in G1/G0, but a significant number of cells are in S- and G2/M-phases. At this level of analysis, it is impossible to distinguish between cells in G0 and G1. However, EdU labeling of DMECs in full growth medium shows that within the length of the typical DMEC cell cycle (~28 hours), nearly all cells pass through S-phase suggesting that there are no cells in G0 in growth medium (not shown). In BPM, we see a nearly complete loss of cells in S and G2/M and a concomitant increase in cells (>91%) arrested in G0/G1. As we will show later in the section titled “Features of Growth Factor-Induced Population Dynamics,” we have reason to believe that the dominant *peak* in BPM (Fig 1C, right panel) contains cells in G0. Therefore, BPM offers desirable dynamic range, reduces background pro-proliferative signaling, and isolates DMECs in G0 or a quiescent state.

The realization that DMECs not only cannot proliferate in low growth factor conditions but in fact die led us to wonder if the overall low proliferation rate in BPM was achieved through a balance of birth and death (see Box 1). To measure dead cells, we optimized the use of the cell-impermeable DNA-binding dye, YOYO1. With this dye, a dead cell can be easily seen as the mCherry signal fades, membrane integrity is lost, cellular DNA is exposed, and the YOYO1 signal appears (see Fig. 1D). The dye is easily detected at low concentrations, stable throughout the timeframe of the assay (not shown) and appears to be passive in that it has no effect on proliferation (Fig S1F). Importantly, cell death in BPM was infrequent and occurred in a reproducible manner with a cell rounding up and blebbing, followed by death with little further fragmentation and the cell most often remaining attached in place. The exception to remaining attached was an occasional tendency for living, motile cells to adhere to the dead cell body. These favorable features allowed for the quantitation of cell death using YOYO-1.

In BPM, there was a small but reproducible death of cells (Fig. 1E) revealing that the low level of net proliferation in BPM is due to a balance between the birth and death rate. Since DMECs are a primary cell isolate rather than a transformed cell line, we assume them to be more sensitive to growth factor levels and more susceptible to pro-apoptotic signals. Give the morphological changes seen during cell death (Figure 1D), we hypothesized that the death in BPM was due to apoptosis. The spontaneous cell death in BPM appears to be apoptosis as it can be inhibited by pretreatment with the caspase inhibitor, Q-VD-OPh (Fig. 1F, S1G). As a negative control, pretreatment with necroptosis inhibitor necrostatin was ineffective at blocking cell death in BPM (Fig. S1H).

In summary, we have generated an experimental system that allows for the quantitative study of *in vitro* endothelial cell proliferation. We can accurately count living and dead cells accurately without considerable impact on proliferative behavior. BPM reduces background pro-proliferative signaling, offers significant dynamic range and models the cell cycle state of endothelial cells in resting blood vessels. Furthermore, BPM creates a state where birth and death are balanced and the effect that growth factors may have on this balance can be examined.

#### Identification of Growth Factors with Pro-Proliferative Behavior in the Experimental System

Having established an experimental system for studying the proliferation of endothelial cells, we sought to identify growth factors with pro-proliferative activity. Through literature search we identified more than 30 growth factors suggested to be pro-angiogenic. We obtained 33 from commercial sources and tested them in the DMEC proliferative setup described above (Table S1). We were disappointed to see that the vast majority of these reported growth factors had no effect on DMEC proliferation. The source of discrepancy between reported results and our results is not immediately clear but may include differences proliferation assays as well as in endothelial cell isolates. We found that four families of growth factors were capable of increasing proliferation: members of the FGF (FGF1 and 2), VEGF (VEGFA165 and VEGFA145), and EGF (EGF and TGFα) families along with HGF. The effects of EGF were significantly weaker than the others and therefore was not included in subsequent analysis. It must be noted that both FGF and HGF are commonly indicated as sources of resistance to VEGF blockade in many cancers, including RCC (Mollica et al. 2019; Zhou et al. 2016).

#### Features of Growth Factor-Induced Population Dynamics

Having generated and characterized an experimental system to study the proliferative effects of identified growth factors on DMECs, we sought to determine mathematical definitions for proliferation, birth, and death rates. Although the proliferation rate in full growth medium was constant over time, it was not immediately clear that the proliferation rate in response to starvation and stimulation with single purified growth factors would behave similarly. The data for each well is quite noisy and the detailed study of the proliferation, birth, and death rates requires numerical differentiation of the noisy data. To limit the effect of applying noise-amplifying numerical process to noisy data, we performed a set of experiments with hundreds of replicates (wells in a 384 well plate) for each condition and applying data smoothing where necessary to generate easily interpretable data. As it is appreciated that such smoothing can affect the exact timing of events, here we focus on a qualitative description of the data and analyses with the intention of using it to arrive at simple definitions for the rates of interest that do not require numerical differentiation.

The number of cells per well (*N*) in response to addition of purified FGF2 is shown as a function of time in Figure 2A. In contrast to the simple case of exponential growth in full growth medium (Fig. 1B), there appears to be two phases: slow steady population growth from the initiation of the experiment to around 24 hours followed by a second phase of more rapid proliferation. The first phase displays mild dependence on growth factor concentration while the second phase showed strong dose dependence. At this level it is not clear whether the time of entry into the second phase is sensitive to FGF2 concentration. We note that the duration of the first phase of population growth is consistent with the time it typically takes for cells to exit G0 and enter the cell cycle.

**Figure 2.**
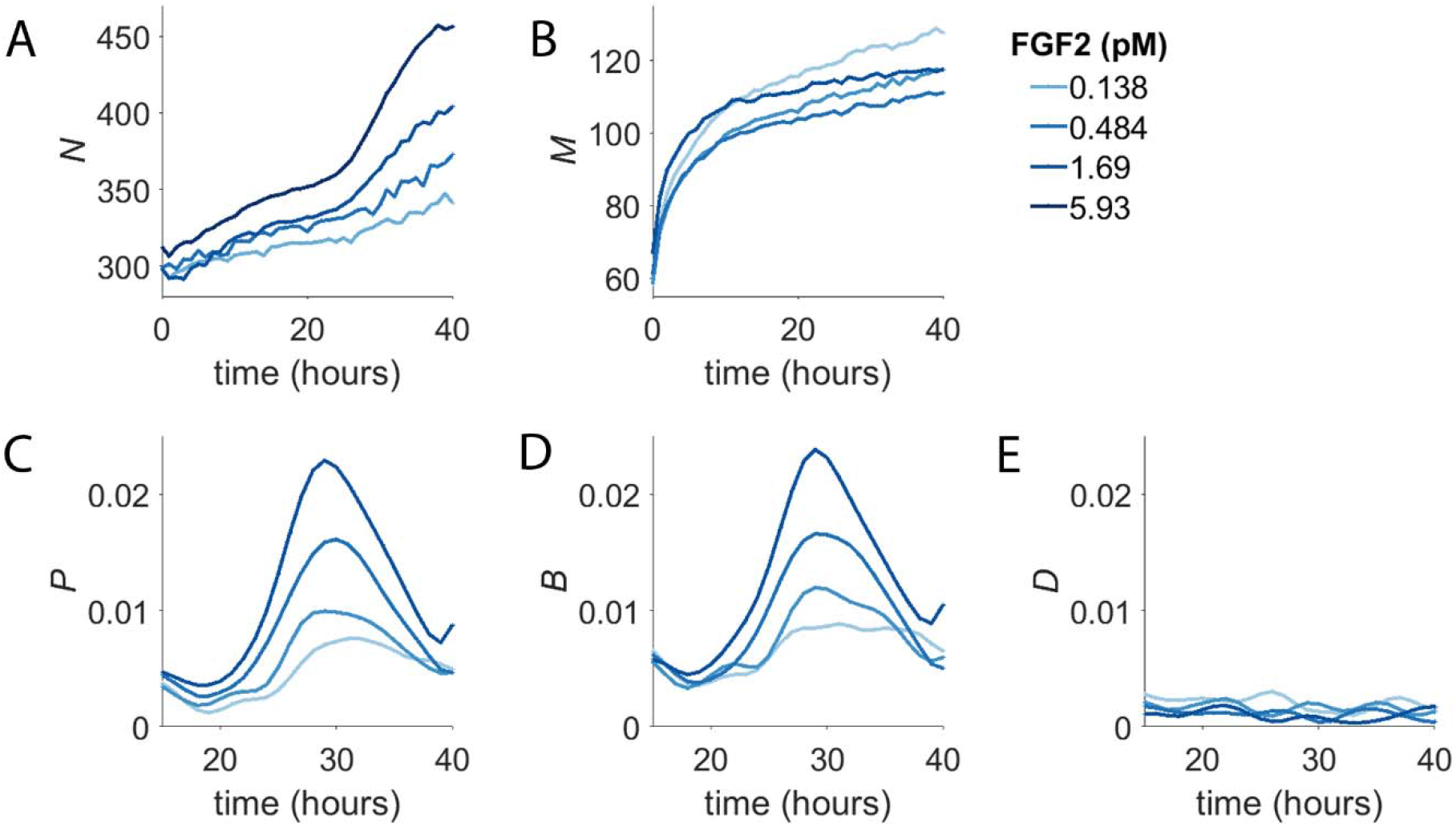
Concentration-Dependent Population Dynamics of Dermal Microvascular Endothelial Cells (DMECs) in the Presence of Pro-Angiogenic Growth Factors. A. The number of cells (*N*) over time for four concentrations of FGF2. B. The number of dead cells (*M*) over time for the experiment shown in panel A. C. The proliferation rate, *P,* for the same experiment as in panels A,B. D. The birth rate, *B,* as a function of time. E. The death rate, *D,* as a function of time.

The number of dead cells per well (*M*) from the same experiment is shown in Fig 2B. It too shows two phases, albeit on a separate time scale compared to the two phases of net population growth seen in Figure 2A. For dead cells, the first phase is short lived transitioning into the second phase after only ~10 hours. Given that the death that is occurs in BPM in the absence of added growth factors is apoptosis (Fig. 1F), the change in death rate over the first few hours following growth factor addition likely reflects the rapid suppression of apoptosis by the presence of growth factor signaling. This suggests that the first phase of population growth seen in Figure. 2A results from the rapid reduction in the basal death rate.

The plots of *N* and *M* over time reveal intriguing characteristics of the concentration-dependent proliferative behavior of DMECs. Taken alone, however, they do not provide deep insight into the proliferative behavior we isolate in the experimental system developed here. To better understand the proliferative behavior of DMECs following the addition of purified growth factors to BPM, we compute the proliferation, birth, and death rates (see Box 1). In general, the rates of interest could be a function of the density of cells (here called *N)* or the time, *t.*

To examine the potential time and density dependence of the endothelial cell proliferation, we computed the proliferation rate and plotted it vs density and time (Fig. 2C). In contrast to density, where we saw no obvious relationship, there is strong time dependence with the proliferation rate peaking at around 30 hours after the addition of growth factor. The rise and fall of the proliferation rate occurred in a dose-dependent manner and the time of maximum proliferation rate was consistent. It begins around the same time, peaks at the same time, and ends at the same time regardless of growth factor dose. The proliferation rate begins to rise around 20 hours and peaks around 30 hours and returns to low level by 40 hours. The mean cell cycle length of these cells was determined by measurement of the doubling time in complete medium to be 21 hours. We interpret the extra time needed to divide during the first division post growth factor treatment to be the passage of cells from G0 in to G1. Thus, the time dependence of the proliferation rate is a direct consequence of the synchronization of the population in G0.

We suspected that the growth factor dependence of the first phase to be the result of growth factors’ rapid effect suppressing apoptosis. Furthermore, the abrupt transition from the first slow phase to the second faster phase was due to the degree of synchrony in the cell cycle introduced by BPM and the time required for cells to re-enter the cell cycle, synthesize DNA, and undergo mitosis. Examination of the birth rate over time (Fig. 2D) showed that, indeed, cells appeared to divide with some degree of synchrony, producing a spike in the birth rate. Meanwhile, the death rate, after the initial rapid decline, proved to be more-or-less constant over the course of the experiment (Fig. 2E). From here, we considered extracting parameters such as the maximum time-dependent proliferation rate or the time of maximum proliferation rate. However, these parameters required more replicates than feasible, given our intention of screen numerous inhibitors in multiple growth factors. For the sake of simplicity in application and interpretation, we applied an exponential model for proliferation. Even though the model assumes the proliferation rate to constant over time, the model fit the raw data reasonably well (Fig. S1I) and resultant proliferation rates were dose-dependent and reproducible.

#### Dose-Response Behavior to Pro-Angiogenic Growth-Factors

The dose-response behavior of pro-angiogenic growth factors is shown in Figure 3. The proliferation rate (Fig. 3A) shows FGF2 to be the most potent and efficacious at driving proliferation. VEGFA and HGF both shared similar efficacy but VEGFA was slightly more potent in that it could produce measurable effects at lower molar concentrations. Since the proliferation rate is the difference between the birth and the death rate (see Box 1) the proliferation rate alone does not completely characterize the system. Comparing the proliferation rate (Fig. 3A) to the birth and death rates (Fig. 3B and C, respectively) shows a more complete story. Two interesting and related features appear: 1.) given the relatively low magnitudes of the death rates the proliferation rate is mostly governed by the birth rate and 2.) growth factors, even at low concentrations push the low death rate closer to zero. Thus, the action of growth factors in the assay is two-fold: first they quickly reduce the death rate while initiating the re-entry into the cell cycle leading to a semi-synchronous wave of birth.

**Figure 3.**
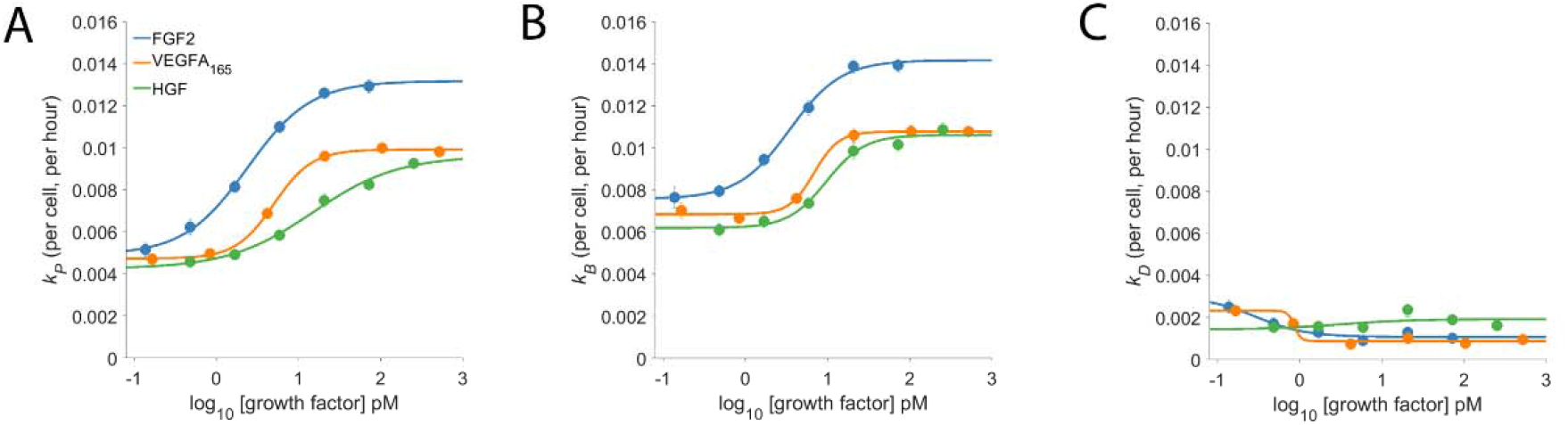
The dose-response proliferative behavior of Dermal Microvascular Endothelial Cells (DMECs) to Three Pro-Angiogenic Growth Factors. The proliferation rate **(A),** the birth rate **(B),** and the death rate (C) for FGF2, VEGFA, and HGF over a range of concentrations. Note that the y-axes share the same scale to facilitate comparison of the relative magnitude of each.

#### Testing a Panel of Kinase Inhibitors

To identify kinases that are used by endothelial cells regardless of the growth factor input, we applied KIR (Rata et al. 2020; Taranjit Singh Gujral, Peshkin, and Kirschner 2014; Taranjit S. Gujral et al. 2014). KIR consists of two steps, first measuring the quantitative effects of 58 kinase inhibitors on proliferation rate in the presence of FGF2, VEGFA, and HGF. The kinase inhibitors used here have been fully characterized against 369 human protein kinases in *in vitro* biochemical assays and have been chosen to provide coverage of the human kinome. The second step compares the biochemical kinase inhibition data for each inhibitor at each dose tested with the degree of inhibition of proliferation within a regularized regression framework to identify kinases important for proliferation.

It was qualitatively apparent that some kinase inhibitors, especially at higher concentrations, produced cell death to an extent that exceeded the apoptosis seen in BPM. Examination of the calculated death rates in the presence of kinase inhibitors confirmed that there were in some cases rapid and massive death. In many cases, the proliferation rate was negative, indicating net loss of cells. These scenarios, where it can be assumed that the birth rate is negligible, provide two independent approximations of cell death. Often in these situations, the loss of cells indicated by the loss of nuclei exceeded the independently measured increased number of dead cells measured via YOYO-1, resulting in unrealistic negative birth rates. This indicates a failure of YOYO-1 to accurately reflect cell death in cases of rapid and extensive death.YOYO-1 was optimized for the detection of death in BPM where the death rate is low and exactly how the error in death rate measured by YOYO-1 increases with increased death rate remains unclear, for the remaining analysis we focus on the proliferation rate which is obtained through high confidence counting of labeled nuclei.

#### Regularization Regression to Identify Universal Kinases

Kinases were identified as the output of regularized regression using the package glmSparseNet (Veríssimo et al. 2018) with the regularization weight alpha set to 0.15 and the kinases were taken as the set of kinases that had nonzero coefficients at the value of lambda with lowest mean-squared error (Fig. S2B). A dataset of mRNA expression levels in DMECs (Davis et al. 2018; ENCODE Project Consortium 2012) was used to identify the set of kinases expressed (Fig. S2A). This procedure produced a ranked order list of kinases important for proliferation for each of the three growth factors tested. The output is shown in Figure 4A. Most kinases appeared in more than one growth factor indicating a degree of similarity in signaling pathways that drive proliferation in endothelial cells. To reduce the complexity of returned kinases and focus on a subset that might prove an enticing target for a more complete antiangiogenic blockade, we chose to compile the intersection of the ranked kinases from each growth factor (Figure 4B). The identity of the kinases in the intersection, ranked by the sum of each kinases’ rank in all three growth factors, are listed in Figure 4C. The kinases in the intersection proved to be a mix of kinases with well appreciated roles in basal metabolism (IR) or proliferation (AKT2 (Shiojima Ichiro and Walsh Kenneth 2002), CDK6 (Sherr, Beach, and Shapiro 2016), PKACG (Yang et al. 2013), and PEAK1 (Wang et al. 2018)) but also some with understudied roles in proliferation (e.g., DCLK2, GRK5/6, and DMPK). Interestingly, the magnitude of the regression coefficient for kinases showed no relationship with the mRNA expression level, suggesting that the method is not simply isolating highly expressed kinases (Fig S3).

**Figure 4.**
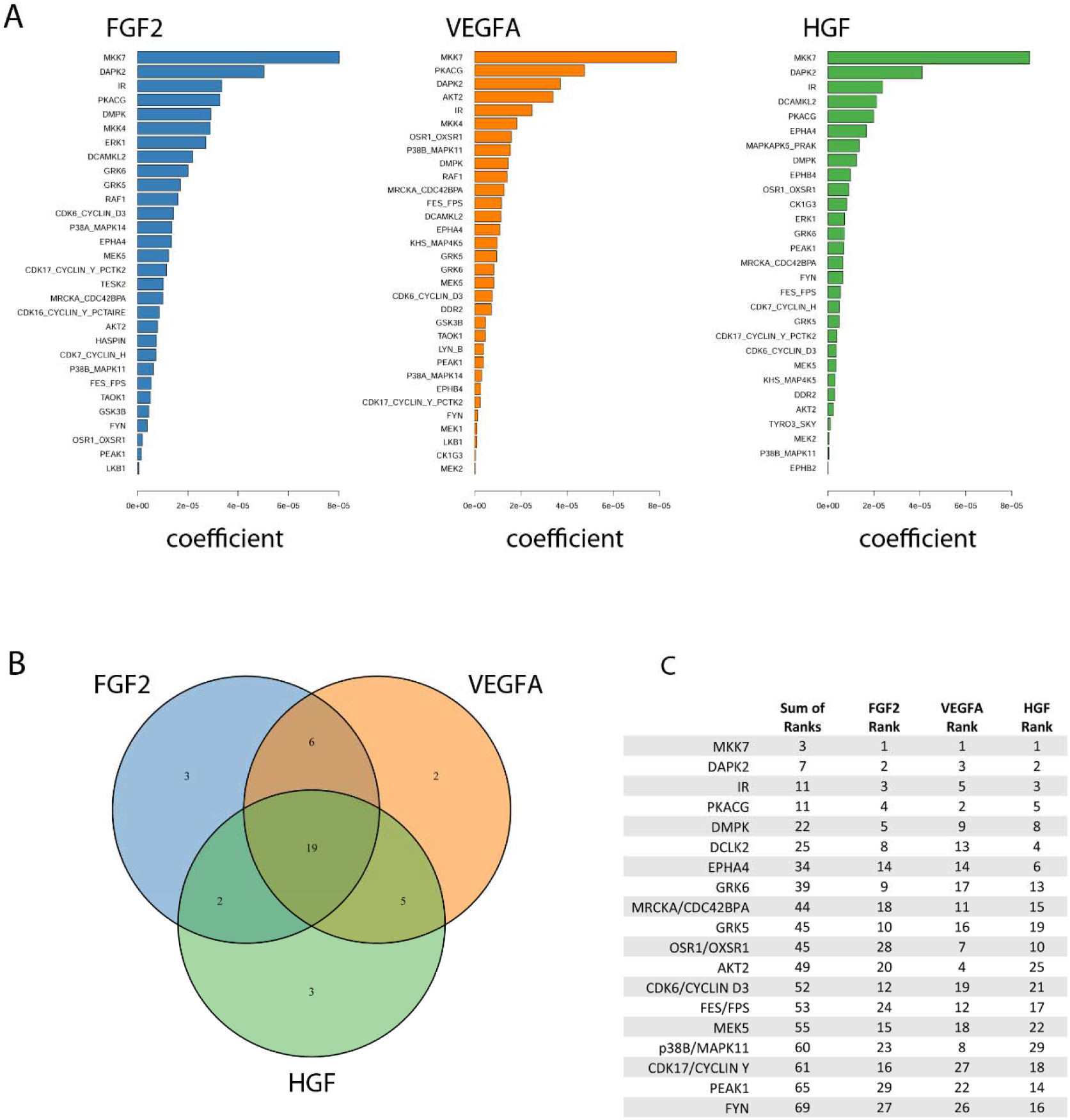
Identification of kinases important for the proliferation rate of Dermal Microvascular Endothelial Cells (DMECs) in each pro-angiogenic growth factor studied. A. Magnitudes of the Kinome Regression (KIR) coefficients (i.e., influential kinases) for each identified growth factor. B. Venn diagram revealing the exclusivity and commonalities in the set of kinases identified for each growth factor. C. The list of the intersection of kinases implicated in all three growth factors. Kinases are ordered according to a rank obtained by summing each kinase’s ranks across all growth factors.

#### Orthogonal Experimental Evidence for Kinases Returned from KIR

We used siRNA to obtain orthogonal evidence that the kinases that were contained within the intersection had impact on the proliferation rate. First, siRNA transfection was optimized for DMECs and the specific plating conditions of the proliferation assay (not shown). We are aware that siRNAs have considerable limitations in this setting for at least two reasons. The first is general to siRNA and is that application of too much siRNA can produce significant off-target effects and therefore false positives while too little might obscure real contributions to proliferation and lead to false negatives. The second is more specific to this system – as large numbers of cells are not easily obtainable from primary DMEC isolates with low replicative potential, independent confirmation of siRNA-mediated knockdown is not feasible. Although, determination of the degree of siRNA knockdown for each kinase of interest could help to reduce false negatives it would not provide any information on false positives. Overall, we wish to address these limitations using a conservative approach, with the goal of leveraging the assay’s reproducibility and quantitative sensitivity to small reductions in proliferation to reduce off target effects at the potential expense of on-target effects.

For the first limitation, we determined the concentration at which siRNAs targeted to each growth factor receptor inhibited the proliferation of DMECs in the presence of its cognate growth factor but not the others. Although there is only one known receptor for HGF (namely MET), the fact that there are four potential FGFRs and three VEGFRs required us to first determine experimentally which receptors are expressed. Via western blot, we identified the receptors expressed to be FGFR1 for FGF2, VEGFR2 for VEGFA, and Met for HGF (Figure S3). We attempted to account for the second limitation by using three siRNAs directed towards each gene of interest and considered the effect to be an average of all three siRNAs.

Using the optimized transfection conditions, we determined the concentration of siRNAs directed toward the expressed growth factor receptors which resulted in a reduction of the proliferation rate in the presence of the cognate growth factor but not the other two. For example, a specific knockdown of FGFR1 would be expected to reduce the proliferation rate of DMECs in the presence of FGF2 but not VEGFA. We found that using 5 nM produced little to no effect while 15 nM produced strong off-target effects (not shown). siRNAs directed toward growth factor receptors at 10 nM appeared to generate specific, on-target reductions in proliferation rate while avoiding off-target responses (Figure 5A). Note that for each target, the reductions in the proliferation rate are incomplete, suggesting that we can indeed measure difference using low concentration of siRNA which should limit non-specific, off-target effects.

**Figure 5.**
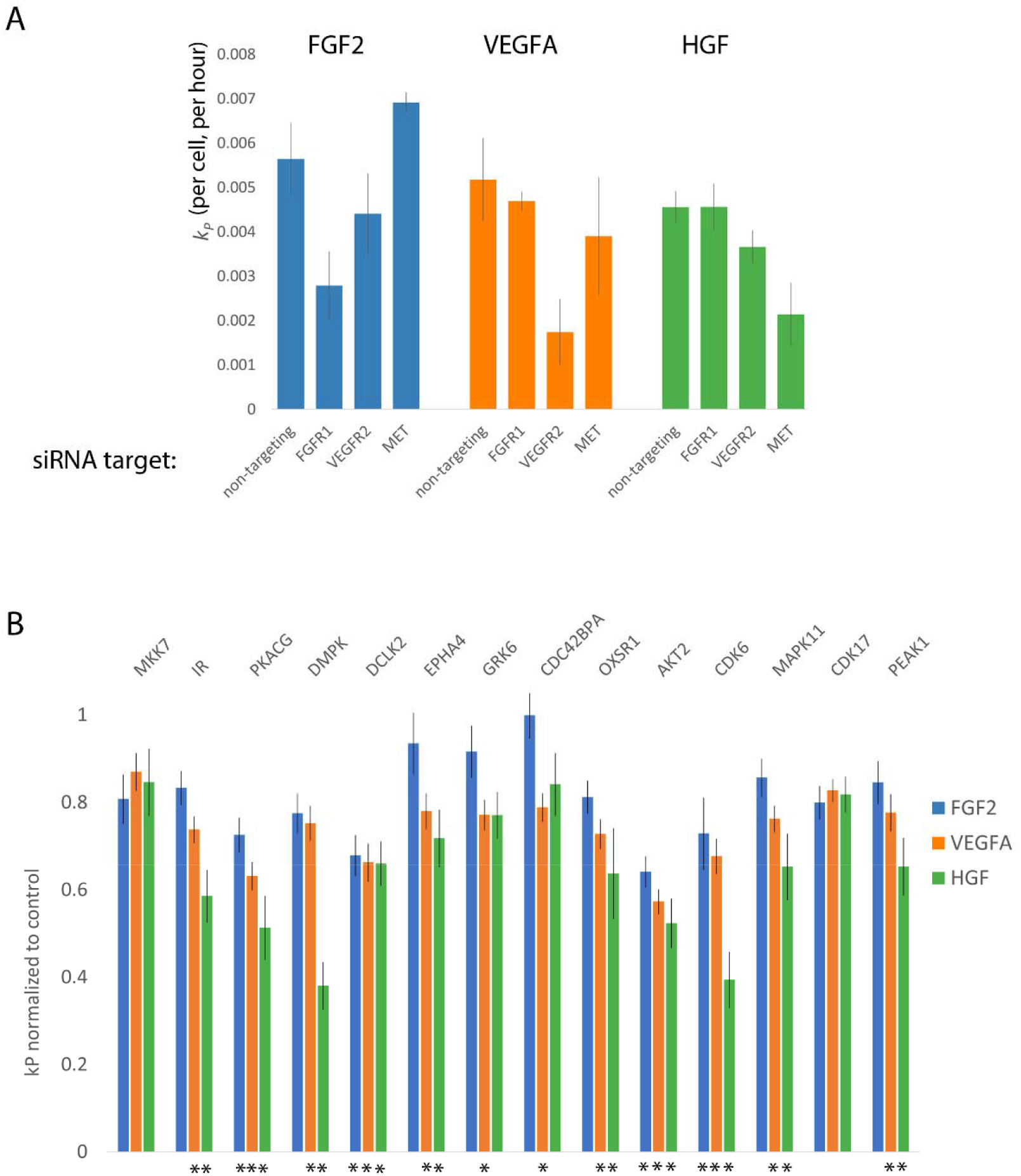
Orthogonal evidence supporting the role of kinases identified by Kinome Regression (KIR) Analysis. A. Targeting growth factor receptors with 5 nM of siRNA reduces the proliferation rate of Dermal Microvascular Endothelial Cells (DMECs) in the presence of cognate growth factors while minimizing reduction in proliferation in other growth factors. B. Targeting kinases with siRNA largely agrees with kinases implicated by KIR. Here, the effect is defined as the average over all three siRNAs used for each kinase and normalized to the no-siRNA control. Error bars are s.e.m. The asterisks indicate growth factors context in which the siRNA targeting a given kinase produced a statistically significant result (*p* < 0.05 from multiple comparison testing).

Next, we tested the intersection kinases using three independent siRNAs to each kinase and the effect of knockdown of each gene was taken to be the average of the effect of all three siRNAs. This conservative approach was chosen because of the difficulty in ascertaining the extent of knockdown in limiting numbers of primary cells on the scale necessary for this work. We present the magnitude of each kinase knockdown in Figure 5B as the proliferation rate following siRNA treatment normalized to the no siRNA control. The magnitude of each effect was tested with ANOVA followed by multiple comparisons. In general, that the siRNA as used resulted in relatively mild to moderate, yet reproducible, inhibition of proliferation. The observed degree of inhibition is consistent with our intention of minimizing off-target responses. All the kinases assessed by siRNA knockdown, except for CDK17 and MKK7 produced a significant reduction in *k_p_* for at least one growth factor. We also note that inhibition of several kinases via siRNA resulted in significant reduction in proliferation in both VEGFA and HGF but not in the most potent driver of proliferation, FGF2. As we are interested in kinases that are involved in the signaling downstream of all three growth factors, we elected to focus on kinases with a statistically significant effect in all three growth factors. The results provide secondary evidence for four kinases to be important for endothelial cell proliferation in FGF2, VEGFA, and HGF: AKT2, CDK6, CAMKL2, and PKACG. Of these, three (AKT2, CDK6, and PKACG) are well characterized. The role of the fourth kinase, DCLK2, in endothelial cell proliferation has not been studied and is therefore of considerable interest as a new regulator of endothelial cell proliferation and angiogenesis.

## Discussion

One of the most appealing aspects of anti-angiogenic therapy was its potential to treat a wide variety of cancers. Unfortunately, current usage of anti-angiogenic therapy remains limited to a smaller subset of cancers than originally envisioned and the efficacy remains lower than hoped. The reason for this limited efficacy remains unclear although redundancy in pro-angiogenic mechanisms is often offered as an explanation. Nonetheless, the fundamental theory behind anti-angiogenic therapy, that all tissues, whether normal or cancerous require adequate blood supply, remains sound. Furthermore, given the fact that angiogenesis is critically involved other pathological disorders as well as in wound healing, the desire for a more thorough understanding of the regulation of the process remains high. Here, we present a novel approach for the identification of critical kinases downstream of growth factor receptors.

There have been many successful efforts to quantify cell proliferation and related analyses have produced many insights into mechanisms governing cell proliferation(Harris et al. 2016; Gross and Rotwein 2016). In general, most approaches fall within two extremes, high through put experiments which tend to rely on endpoint measurements and time-lapse, single cell approaches. High-throughput, end point assays can provide a wealth of data on the effects of an enormous number of perturbations on cell proliferation. An end-point assay may reveal an increase or decrease in the number of cells upon perturbation but offers no insight on whether those changes resulted from impacting birth and/or death. Conversely, detailed single cell tracking-based data can offer information on cell cycle transitions, birth, pro-apoptotic signaling events and death. Despite the development of many tools to analyze such detailed experiments, the throughput remains low. With our assay design, we have tried to thread the needle between the wealth of data provided by increased throughput while also attempting to acquire kinetic data that can provide insight into time-dependent responses.

We observed some interesting aspects of the effects of growth factors on the cell cycle and proliferation. Fig. 2D clearly shows that there is a ‘wave” of cell division upon addition of growth factor that lasts roughly 20 hours. From the raw cell counts in Fig. 2A, we can read off the number of new cells and see that at the highest FGF2 concentrations the population goes from ~320 to 450. In other words, over the course of the growth factor driven wave of birth, only roughly 130 new cells are produced from 320. (Note that this analysis ignores the low level of cell death.) Clearly, the entire population is not dividing upon the addition of FGF, the single most potent and efficacious growth factor in the assay. We see similar behavior for HGF and VEGF (data not shown). Furthermore, the number of cells in that do divide is concentration dependent suggesting that increasing concentrations of growth factors increase the fraction of cells that re-enter the cell cycle. This type of threshold behavior has been seen before in fibroblasts (Gross and Rotwein 2016).

This raises two questions: 1.) What causes a subset of cells to re-enter the cell cycle and, more importantly for the current work, 2.) are the cells that do not re-enter the cell cycle even capable of doing so or is there something about BPM that causes them to enter an irreversible quiescence? All we can definitively say about it is that the DMECs under study were isolated from a single donor to reduce genetic heterogeneity and, more importantly, that we have evidence that the addition of a combination of purified growth factors results in >99% of DMECs plated in BPM for 24 hours dividing (not shown). Thus, the state is not irreversible and results in no artifact vis a vis our analysis of pro-angiogenic kinases. We believe the kinetic assay of proliferation and apoptosis developed here provides a method for addressing this and other unanswered questions regarding these two critical cellular functions and their interaction.

Using KIR, we identified 19 kinases to be important for endothelial cell proliferation in the presence of multiple bona fide pro-angiogenic factors. Furthermore, we also provide additional evidence that four of these 19 are critical for endothelial cell proliferation using an orthogonal method, siRNA-mediated knockdown. At this scale, the siRNA approach has several limitations, including a lack of validation of efficient gene knockdown. To overcome this weakness, we chose to take a conservative approach to out interpretation of the results by focusing on the kinases that were found to statistically impact proliferation while avoiding statements about those that had no large, reproducible effect. Our siRNA results show that within the optimized format, the siRNAs produced mild to moderate effects. Therefore we emphasize that while the knockdown of a kinase producing an effect in all three growth factors certainly provides strong evidence for a role in proliferation, we cannot with great confidence exclude the possibility that those that did result in an effect in one or two growth factors (or even those that had no significant effect at all) do not also have a considerable impact on endothelial cell proliferation. For example, it would seem unlikely that the insulin receptor is somehow not important in FGF driven proliferation. It is interesting that MKK7, which has been implied in pro-angiogenic signaling (Mitchell et al. 2006), dominated the KIR output yet was not corroborated by siRNA. This possibly reflects the differences in between the two methods and their sensitivity to functional redundancy. For example, consider a scenario where two kinases are similar enough to functionally compensate for each other and they share similar sensitivity to a subset of KIR compounds. In this scenario, KIR might return one or both of these kinases. However, siRNAs with specificity to only one of these kinases would fail to produce any effect due to redundancy.

One of these four kinases identified and further validated here, CDK6, has an inhibitor palbociclib approved for treatment of hormone positive, HER2 negative metastatic breast cancer. Indeed, some have speculated that at least some of Palbociclib’s efficacy is due to an anti-angiogenic effect (Liu, Liu, and Chen 2018; Ehab and Elbaz 2016). Recently, palbociclib in combination with taxanes, was shown to increase apoptosis and reduce HIF-1α in a pre-clinical model of squamous cell lung cancer (Cao et al. 2019). Similarly, the pan-Akt inhibitor capivasertib has been shown to improve outcomes in a variety of Akt1 E17K mutant cancers (Kalinsky et al. 2021). Another Akt inhibitor, ipatasertib, in combination with androgen pathway inhibition, improved progression free survival in prostate cancer (Sweeney et al. 2021). Protein kinase A (PKA) has been shown to be important for directly regulating endothelial cell during angiogenesis (Zhao et al. 2019), and indirectly through tumor associated macrophage-based secretion of VEGF (Na et al. 2020).

The present work also indicates the DCLK2 as a novel regulator of endothelial cell proliferation. DCLK1 and 2 are understudied kinases, with the majority of work being upon DCKL1 and its role in cancer (Ferguson et al. 2020), neuronal survival (Nawabi et al. 2015), and maintenance of intestinal crypt cell stemness (Chandrakesan et al. 2017). DCLK2 was identified as having synthetic lethal interaction with mutant Ras in colorectal cancer (Luo et al. 2009), suggesting that it may be an enticing target for cancers drive by the Ras oncogene. It will be of interest to further explore the role of DCLK2 in endothelial cell biology and cancer.

## Methods

### Growth Factors and Cytokines

All human recombinant growth factors and cytokines were purchased from PeproTech (NJ) and resuspended in the solvent recommended by the manufacturer at 100μg/mL, aliquoted, and frozen until use.

### Cloning

The viral transfer plasmid expression a nuclear-localized mCherry was obtained using standard cloning techniques. First, pLVX-EF1α-mCherry-N1 (puro) was digested with EcoRI and NotI. The 8.5kB band representing the backbone of the vector was excised and purified. The nuclear-localized mCherry was digested from pBRY-nuclear mCherry-IRES-PURO (Addgene plasmid 52409) and the ~860bp band was gel purified. The insert was ligated into the pLVX vector overnight at room temperature, transformed into Stbl3 bacteria and plated on LB/ampicillin plates. Colonies were picked, cultured overnight, and tested for proper ligation using digestion with EcoRI and NotI. Positive clones were tested be sequencing and the sequence of both the 3X-NLS and mCherry was confirmed to be correct.

### Cell Culture, Virus Production, and Viral Transduction

Human Dermal blood microvascular endothelial cells (DMECs) isolated from a single donor were purchased from Lonza (CC-2183). DMECs were cultured in EGM2-MV (Lonza) and passaged using trypsin and trypsin neutralization solution (Lonza). Viruses carrying a gene for the expression of nuclear localized mCherry were produced via triple transfection of HEK-293T cells using FuGENE. The plasmids used were psPAX2, pMDG.2, and pLVX-E1α-NLS-mCherry. Twenty-four hours after transfection, the medium was replaced. For the next 3 days, the supernatant was collected, spun at 500xg to remove large debris, and stored at 4C. After all supernatant was collected, the pooled supernatant was passed through a 0.45 μm filter and transferred to centrifuge and centrifuged for 90 minutes in a SW28 singing bucket rotor. After centrifugation, the supernatant was discarded and the pellet containing concentrated virus was resuspended in 100 μL of PBS. To create a large supply of labeled DMECs, three tubes of purchased DMECs were thawed, counted, plated at 5000 cells/cm^2^ and cultured overnight. The next day purified and concentrated virus was added to the cells at MOI = 5 in medium containing 8 μg/mL polybrene. The next day, the medium was changed, and the cells were cultured continuously, passaged when ~80% confluent until reaching the sixth passage (cells are purchased at passage three). At this point, the cells were trypsinized, pelleted, counted, and resuspended in EGM2-MV supplemented with 10% DMSO at a density of 500,000 cells/mL. The labeled DMECs were aliquoted in 500 μL aliquots and frozen at −80C overnight and then transferred to long term storage in liquid nitrogen. For every experiment presented here, tubes were thawed, viable cells determined and plated at 5000 cells/mL, cultured for 6 days with medium changes before being collected, counted, and plated. For proliferation assay in 384 well plates, cells were plated at 255 cells per well or 3591 cells/cm^2^.

### Formulation of Basal Proliferation Medium (BPM)

BPM contains 0.5% human AB serum (0.5%), insulin, r-Transferrin, 0.5% BSA, hydrocortisone.

### The proliferation assay

YOYO-1 iodide was purchased from ThermoFisher (Y3601) and added to BPM immediately before plating cells for the proliferation assay. DyeCycle Green was purchased from ThermoFisher and used as a live cell stain at a final concentration of 10 μM to count living nuclei and determine the labeling efficiency of nuclear mCherry transduction and expression.

### Image Analysis for Counting Nuclei and Dead Cells

Time-lapse imaging of DMEC proliferation was done in an IncuCyte ZOOM (Sartorius) with a 4x objective. Nuclei and dead cells were counted using the built-in image analysis capabilities of the IncuCyte ZOOM.

### Cell Cycle Distribution and EdU Incorporation

DMECs were plated on poly-L-lysine-coated 6-well glass-bottom plates at 8,450 cells per cm^2^ in either BPM or growth medium (EGM2-MV, Lonza). After 24 hours, cells were fixed with 4% paraformaldehyde in PBS for 10 minutes, permeabilized with 0.1% triton in PBS for 10 minutes, then stained with DAPI (Cell Signaling) for one hour at room temperature. Following staining, cells were washed once with PBS and then imaged immediately on a Nikon at 10x.

For long-term EdU incorporation, we first optimized the concentration of EdU that could be detected but not greatly alter the proliferation rate. We did this by plating cells in varying concentration of EdU in growth medium and fixing after various times. We found that culture in 240 nM EdU for 48 hours resulted in a less than 5 % reduction of the proliferation rate while being incorporated into ~95% of cells. For experiments, we then plated cells in BPM, and 24 hours later, added EdU with the growth factor or added complete growth medium.

### ELISA

Pre-coated ELISA plates were purchased from R&D Systems and protocols were performed according to manufacturer instructions. Cell culture supernatants were obtained by plating 600 cells per well in 384-well plates in 60 μL BPM. After 24 hours, 20 μL of BPM containing growth factors at 4x intended final concentration were added. Twenty-four hours following growth factor addition, 60 μL of supernatant was removed from each well and pooled according to growth factor and concentration. The pooled supernatants were then clarified by centrifugation to remove any cell debris and frozen at −20C until all replicates were performed. Supernatants were thawed to room temperature and utilized immediately in the ELISA.

### Analysis of kinetic measurements of cellular proliferation and death

We applied the following definitions, as approximation to obtaining the proliferation rate from an exponential fit, to calculate per capita proliferation rates for each well:

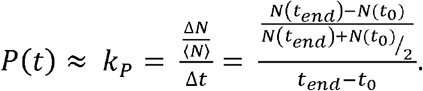

Where *N*(*t*) is the number of nuclei as a function of time, 〈*N*〉 is the average number of nuclei over the experiment, and *t*_0_ and *t_end_* is the first and last time points in the experiment. The death rate was defined similarly:

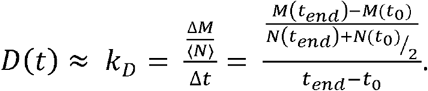

The birthrates were then calculated as *k_B_* = *k_P_* + *k_D_*. These definitions assume that the rates are independent of the overall density and time. Although we show that density is not a critical factor (Figure S1E), the rates are clearly dependent on time (Figure 2). We use these definitions as they are simple to compute and calculation of time-dependent rates were not feasible with the noise inherent in the data.

### Kinase Inhibitor Screen

Extensively characterized kinase inhibitors were tested in the proliferation assay in the presence of each of the three growth factors. Inhibitors were tested with four replicates at each of six concentrations, the highest being 5 μM with serial 3-fold dilution.

### Kinome Regression (KIR)

Kinome regression was performed as regularized regression using the R package glmSparseNet. The input was the calculated difference between each inhibitor/growth factor combination and the appropriate control (DMSO/growth factor combination) contained on the same plate. The kinase data from the in vitro characterization was filtered according to endothelial cell expression (www.encodeproject.org, identifier: ENCFF110UGQ) using a cutoff of 1.5 arbitrary expression units (see Figure S2). The following parameters were used in the analysis: α = 0.15 and standardize = TRUE.

### siRNA Validation of Kinases Identified by KIR

We tested four different siRNA transfection reagents (RNAiMAX, DharmaFECT 4, for two properties: 1.) To efficiently knock-down a gene of interest and 2.) to have minimal impact on proliferation in the assay. To evaluate transfection efficiency, we used KIF11 which is critical for cytokinesis and when knocked down results in easily scored rounded cells. We found that 1uL/mL of RNAiMAX resulted in 95% rounded cells in while decreasing the proliferation rate by less than 3%. We knocked down kinases using siRNAs from Qiagen (see spreadsheet for the ordering numbers of the siRNAs used) at 10 nM. For transfection, cells were plated in BPM containing YOYO1 in 384 well plates as usual for the proliferation assay. While the cells were adhering, siRNA transfection complexes were formed after a 10-minute incubation and added to the cells with four replicates. Twenty-four hours after plating/transfection, growth factors were added at the following concentrations: FGF2, 1.25 ng/mL; VEGFA, 20 ng/mL, and HGF, 20 ng/mL. The plates were then added to the IncuCyte ZOOM and imaged every two hours for 48 hours. The proliferation rates were extracted as described in the section.

## Acknowledgements

This work was funded by the NICHD award HD091846 and NIA award AG073341 to LP and MWK.

## SUPPLEMENTARY FIGURE LEGENDS

**Figure S1.**
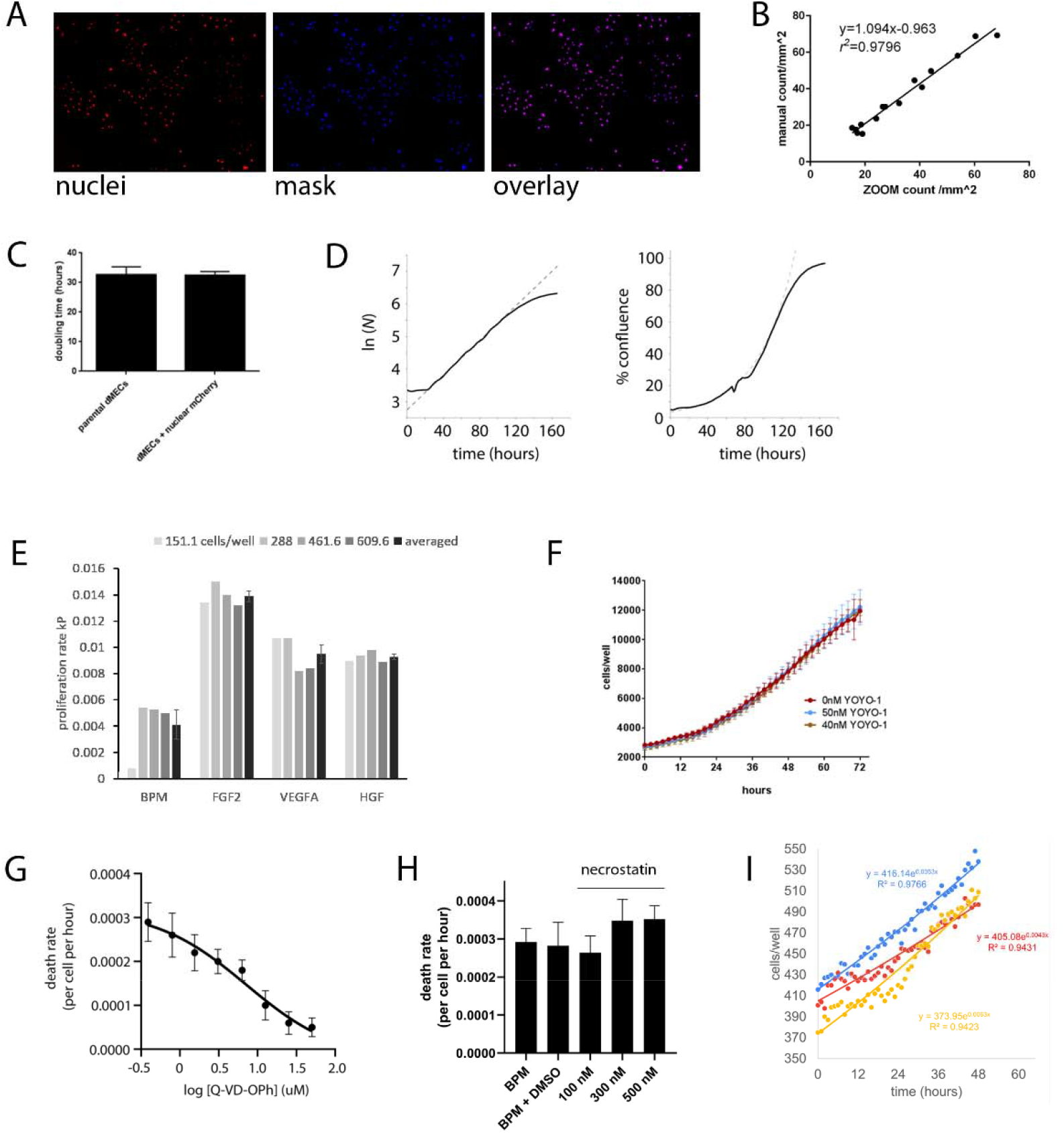
More on the Establishment of the Experimental System. A.) Nuclei can be counted via image processing. The right panel shows a fluorescent micrograph of nuclear mCherry labeled Dermal Microvascular Endothelial Cells (DMCEs), while the center panel shows the mask generated by automated image analysis. The right panel shows the overlay of the two. B.) There is a high degree of correspondence between automated counting (x-axis) and manual counting of nuclei (y-axis). Additionally, the error is apparently independent of density. C.) Labeling DMECs with nuclear mCherry via lentivirus-mediate gene transfer does not alter the doubling time of DMECs. D.) Left panel: Nuclear labeled DMECs produce exponential growth over an interval of time as evidenced by linear relationship between time in hours (x-axis) and the natural logarithm of cell number (y-axis). Right panel: Percent confluence for the same experiment. Note the deviation from exponential growth occurs at ~120 hours or when confluency grows above ~70%. For both panels, solid line is data, dashed line is a fitted exponential curve. E.) The proliferation rate (defined as in main text, section *Features of Growth Factor-Induced Population Dynamics)* is independent over a range of plating densities in the presence of growth factors. F.) YOYO-1, at concentrations as high as 50 nM, have no effect on the proliferation of DMECs. G.) The inhibition of death rate in (Basal Proliferation Medium) BPM (defined as in main text, section *Features of Growth Factor-Induced Population Dynamics)* is dependent on the concentration of inhibitor. H.) The death rate in BPM is not inhibited by relevant concentration of the necroptosis inhibitor necrostatin. I.) The raw data for cells proliferating in response to growth factors added to BPM (here 20 ngs/mL VEGFA) are well approximated by an exponential fit.

**Figure S2.**
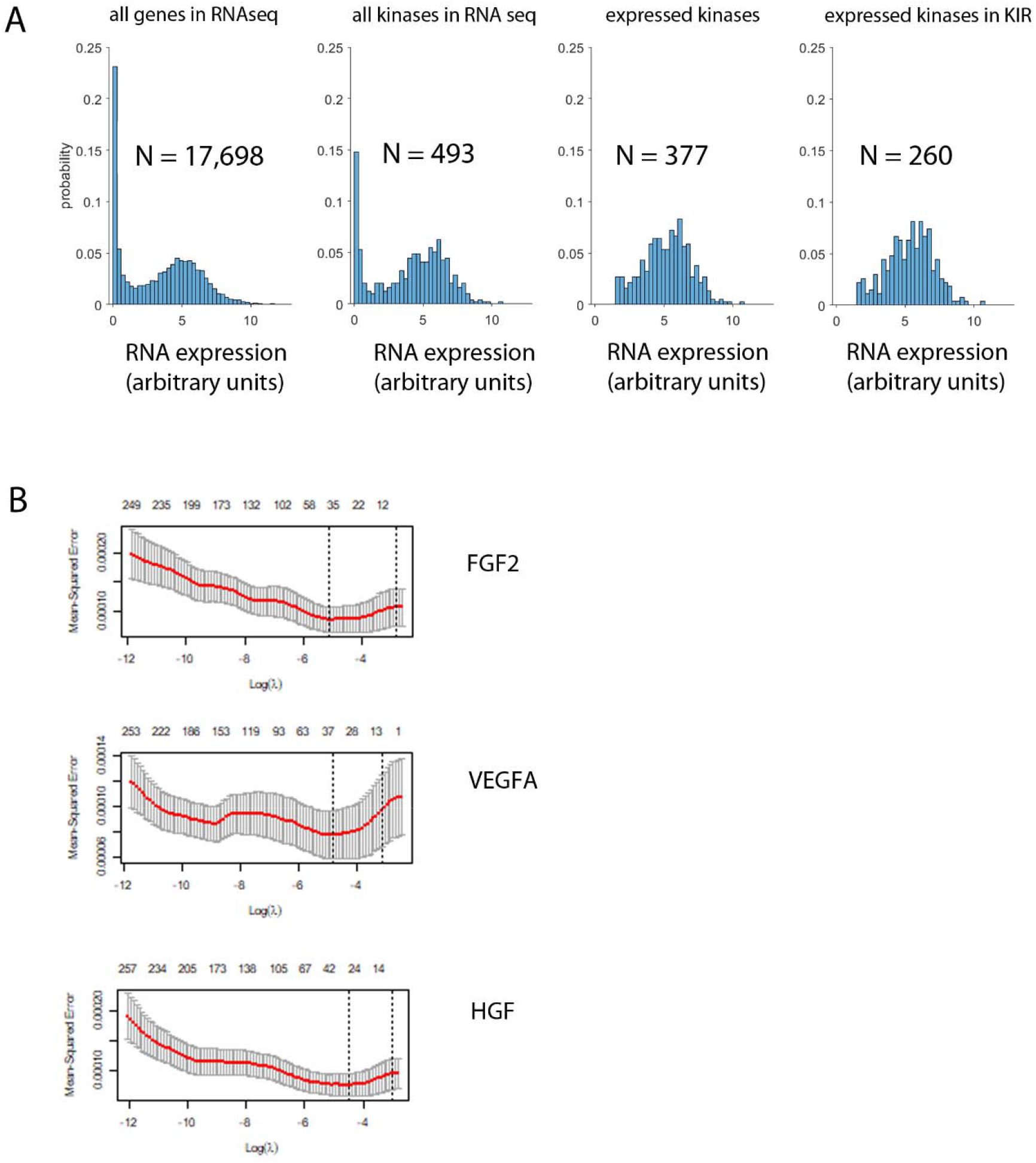
Additional data regarding Kinome Regression (KIR). A.) From left to right we detail the RNA-seq dataset used to identify the set of kinases expressed by Dermal Microvascular Endothelial Cells (DMECs). On the far left is shown the histogram of expression levels of all 17,698 genes in the dataset. The bimodal nature of the expression data is easily seen and suggests that genes in the first peak with relatively low expression values are not actually expressed while those in the second more broad peak with higher expression values are. Moving rightward, the histogram of the expression of only protein kinase genes (*N* = 493) is shown. It can be seen that kinases too follow the expected bimodal distribution as seen for all genes. We use these two observations to justify the selection of a threshold for real kinase expression. This threshold was set at 1.5 arbitrary units. Next histogram shows the distribution of the 377 expressed kinases in DMECs. Finally, on the far right we see the histogram of the intersection between expression kinases and KIR kinases, i.e., those for which the compounds have been thoroughly characterized. Thus, we arrive at 265 kinases which could potentially be returned from KIR analysis. This is not the final number as will be detailed next. Note that in Fig. S2A the number of kinases in the RNA-seq dataset is 493. This is 25 shy of the total number of kinases thought to be in the human genome. Of the 25 kinases not in the expression data, five are KIR kinases. To be conservative, we include those kinases in our analysis. B.) Plots of log(λ) vs mean-squared error (MSE) from leave-one-out cross-validation. We used the kinases from the model that minimize MSE (vertical dashed line).

**Figure S3.**
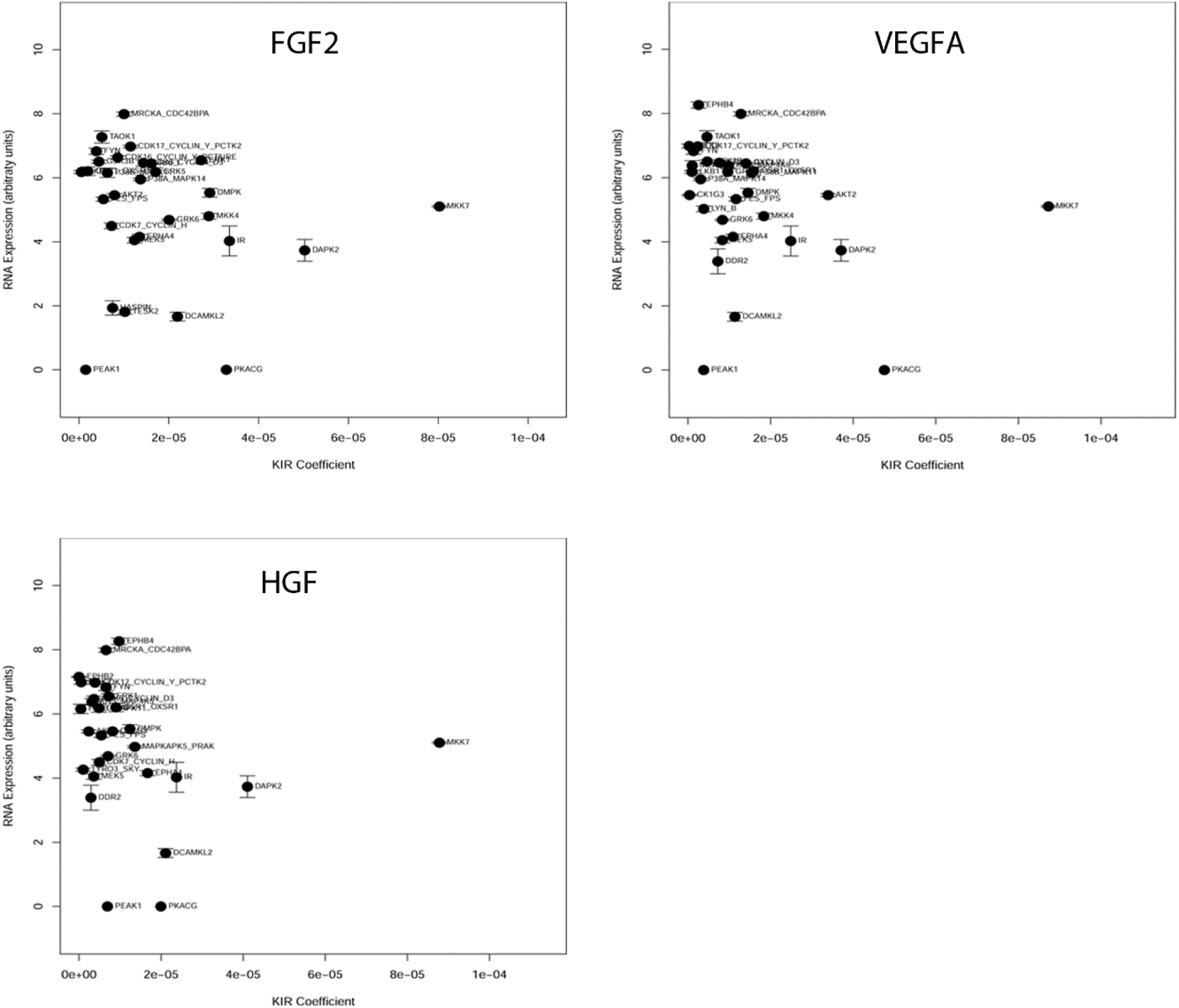
The Kinome Regression (KIR) coefficients plotted against the mRNA expression level. Note the lack of relationship between expression level and magnitude of contribution. It can also be seen that two kinases for which we lack expression data, PEAK1 and PKACG, were returned by the analysis.

**Figure S4.**
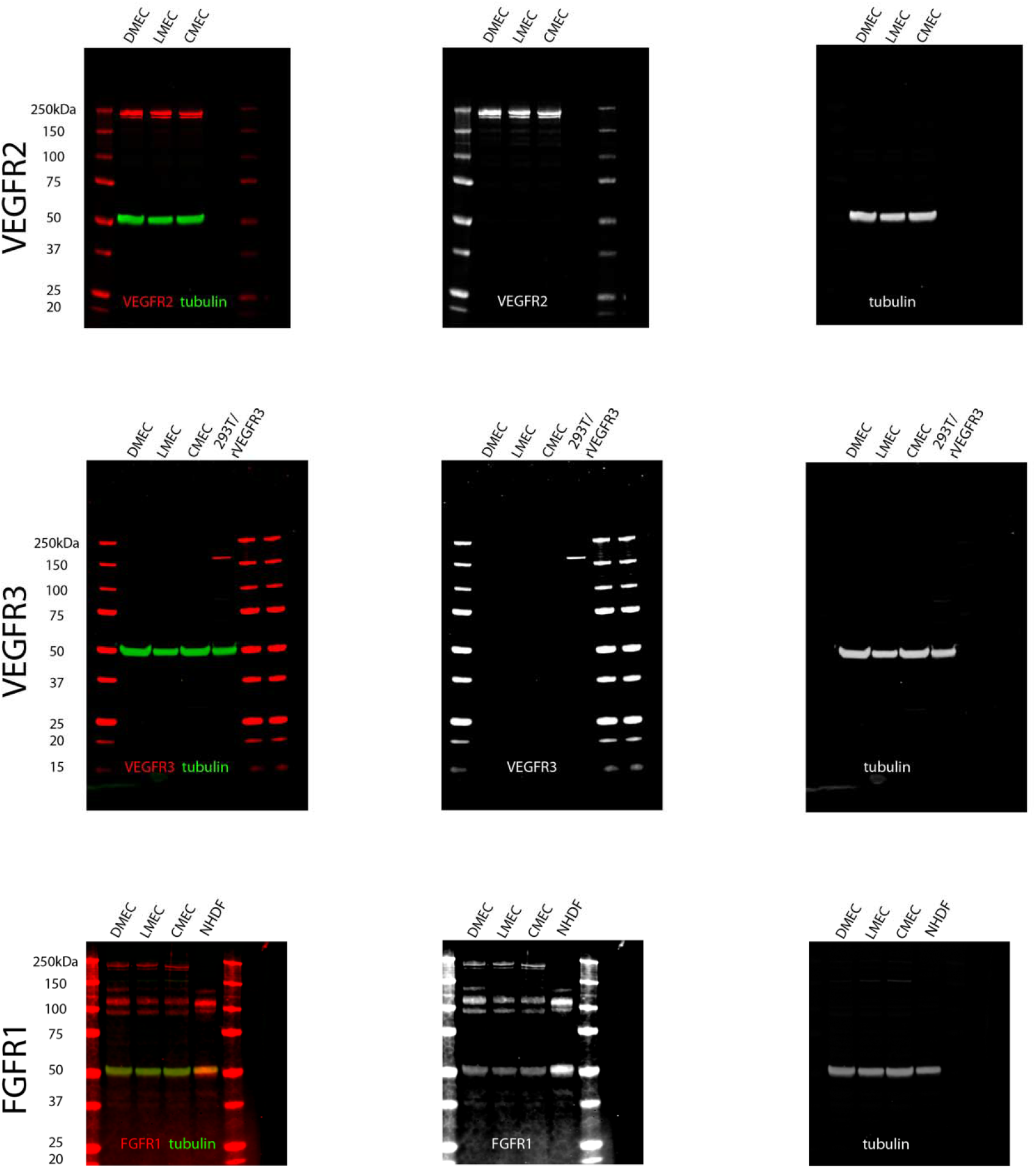
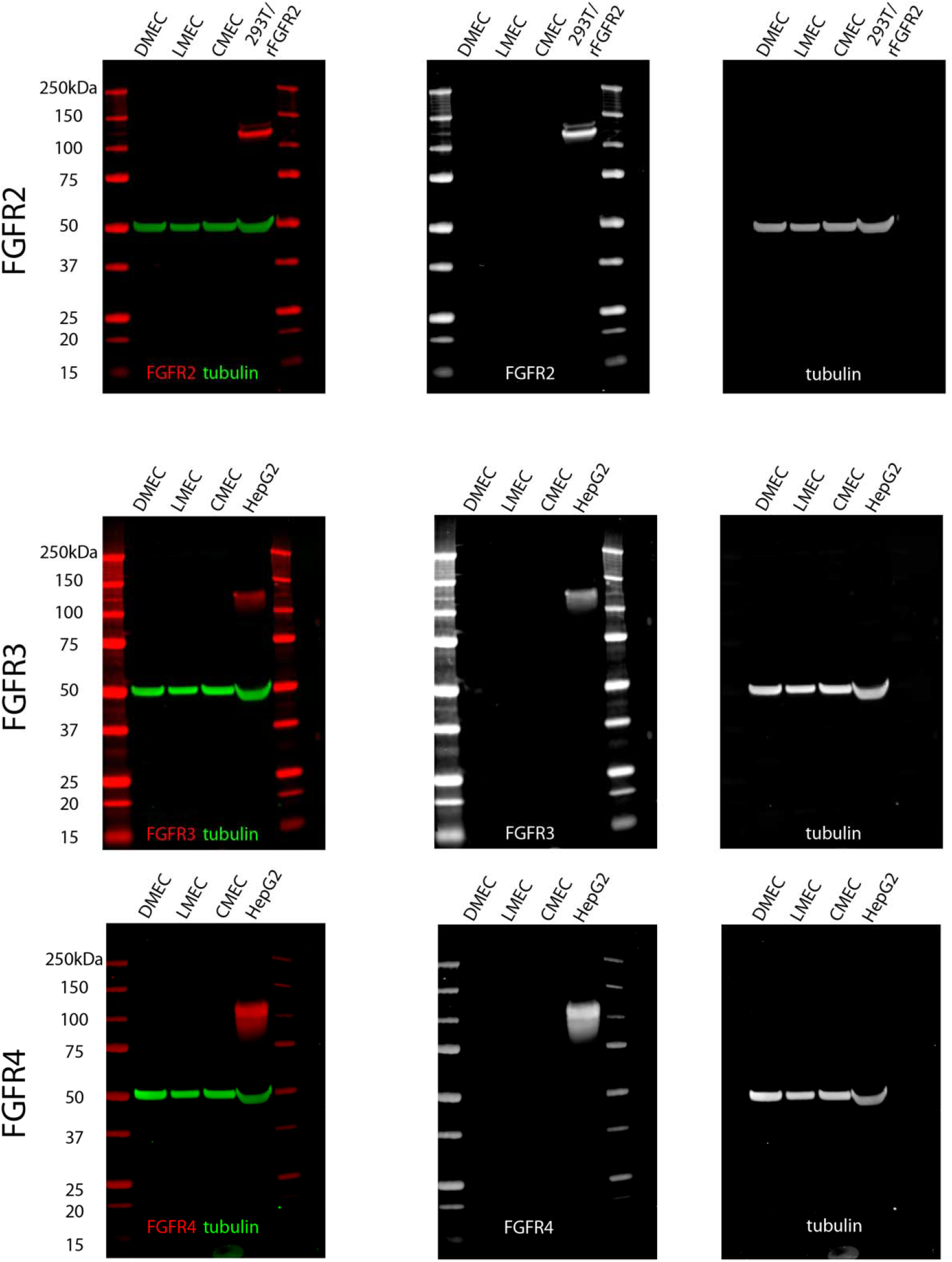
The identification of relevant growth factor receptors expressed on Dermal Microvascular Endothelial Cells (DMECs) via western blotting. There are two VEGF receptors in the genome with kinase activity, VEGFR2 and VEGFR3. DMECs only express VEGFR2. Similarly, there are four FGFRs in the human genome and DMECs only express FGFR1.

**TABLE S1.**

| Growth Factor | Supplier | MW (kDa) | Highest Concentration Tested (nM) | Pro-Proliferative? |
| --- | --- | --- | --- | --- |
| **Activin A** | PeproTech | 26 | 3.85 | N |
| **Ang1** | PeproTech | 56.3 | 1.78 | N |
| **Ang2** | PeproTech | 50.1 | 2 | N |
| **CTGF** | PeproTech | 11 | 9.09 | N |
| **EG-VEGF** | PeproTech | 9.6 | 10.42 | N |
| **EGF** | Millipore | 133.95 | 0.01 | Y (weak) |
| **FGF1** | PeproTech | 16.8 | 5.95 | Y |
| **FGF2** | PeproTech | 17.2 | 5.81 | Y |
| **FGF3** | R&D | 21.1 | 7.11 | N |
| **G-CSF** | PeproTech | 18.7 | 5.35 | N |
| **GM-CSF** | PeproTech | 14.6 | 13.7 | N |
| **Heregulin 1** | PeproTech | 7.5 | 13.33 | N |
| **HGF** | PeproTech | 79.4 | 2.52 | Y |
| **IGF-1** | R&D | 7.6 | 26.32 | N |
| **IL-1β** | R&D | 17.3 | 5.78 | N |
| **IL8** | PeproTech | 8.9 | 11.24 | N |
| **Leptin** | R&D | 16 | 62.5 | N |
| **MCP-1** | PeproTech | 8.6 | 11.63 | N |
| **PDGF-BB** | R&D | 24.3 | 4.12 | N |
| **PIGF** | PeproTech | 29.7 | 5.05 | N |
| **SDF-1α** | PeproTech | 8 | 25 | N |
| **TGF-α** | PeproTech | 5.5 | 18.18 | Y (weak) |
| **TGF-β1** | PeproTech | 25 | 4 | N |
| **TGF-β2** | PeproTech | 25 | 4 | N |
| **TGF-β3** | PeproTech | 25 | 4 | N |
| **VEGFA 121** | R&D | 28 | 5.36 | Y (weak) |
| **VEGFA 145** | R&D | 34 | 4.41 | Y |
| **VEGFA 165** | PeproTech | 38.2 | 3.93 | Y |
| **VEGF B** | PeproTech | 38 | 3.95 | N |
| **VEGF C** | PeproTech | 27 | 5.56 | N |
| **VEGF D** | PeproTech | 26.2 | 5.73 | N |
| **Wnt1** | PeproTech | 38.4 | 2.6 | N |
| **Wnt7B** | PeproTech | 35.5 | 2.82 | N |

## References

Cabral, Thiago, Luiz Guilherme M. Mello, Luiz H. Lima, Júlia Polido, Caio V. Regatieri, Rubens Belfort Jr, and Vinit B. Mahajan. 2017. “Retinal and Choroidal Angiogenesis: A Review of New Targets.” International Journal of Retina and Vitreous 3 (August): 31.

Cao, Joan, Zhou Zhu, Hui Wang, Timothy C. Nichols, Goldie Y. L. Lui, Shibing Deng, Paul A. Rejto, et al. 2019. “Combining CDK4/6 Inhibition with Taxanes Enhances Anti-Tumor Efficacy by Sustained Impairment of PRB-E2F Pathways in Squamous Cell Lung Cancer.” Oncogene 38 (21): 4125–41.

Carmeliet, Peter. 2005. “VEGF as a Key Mediator of Angiogenesis in Cancer.” Oncology 69 Suppl 3 (November): 4–10.

Chandrakesan, Parthasarathy, Jiannan Yao, Dongfeng Qu, Randal May, Nathaniel Weygant, Yang Ge, Naushad Ali, et al. 2017. “Dclk1, a Tumor Stem Cell Marker, Regulates pro-Survival Signaling and Self-Renewal of Intestinal Tumor Cells.” Molecular Cancer 16 (1): 30.

Choueiri, Toni K., Bernard Escudier, Thomas Powles, Paul N. Mainwaring, Brian I. Rini, Frede Donskov, Hans Hammers, et al. 2015. “Cabozantinib versus Everolimus in Advanced Renal-Cell Carcinoma.” The New England Journal of Medicine 373 (19): 1814–23.

Collins, M. K., G. R. Perkins, G. Rodriguez-Tarduchy, M. A. Nieto, and A. López-Rivas. 1994. “Growth Factors as Survival Factors: Regulation of Apoptosis.” BioEssays: News and Reviews in Molecular, Cellular and Developmental Biology 16 (2): 133–38.

Crawford, Talia N., D. Virgil Alfaro 3rd, John B. Kerrison, and Eric P. Jablon. 2009. “Diabetic Retinopathy and Angiogenesis.” Current Diabetes Reviews 5 (1): 8–13.

Davis, Carrie A., Benjamin C. Hitz, Cricket A. Sloan, Esther T. Chan, Jean M. Davidson, Idan Gabdank, Jason A. Hilton, et al. 2018. “The Encyclopedia of DNA Elements (ENCODE): Data Portal Update.” Nucleic Acids Research 46 (D1): D794–801.

Ehab, Moataz, and Mohamad Elbaz. 2016. “Profile of Palbociclib in the Treatment of Metastatic Breast Cancer.” Breast Cancer 8 (May): 83–91.

ENCODE Project Consortium. 2012. “An Integrated Encyclopedia of DNA Elements in the Human Genome.” Nature 489 (7414): 57–74.

Ferguson, Fleur M., Behnam Nabet, Srivatsan Raghavan, Yan Liu, Alan L. Leggett, Miljan Kuljanin, Radha L. Kalekar, et al. 2020. “Discovery of a Selective Inhibitor of Doublecortin like Kinase 1.” Nature Chemical Biology 16 (6): 635–43.

Grépin, Renaud, and Gilles Pagès. 2010. “Molecular Mechanisms of Resistance to Tumour Anti-Angiogenic Strategies.” Journal of Oncology 2010 (March). https://doi.org/10.1155/2010/835680.

Gross, Sean M., and Peter Rotwein. 2016. “Unraveling Growth Factor Signaling and Cell Cycle Progression in Individual Fibroblasts.” The Journal of Biological Chemistry 291 (28): 14628–38.

Grünwald, Viktor, and Axel Stuart Merseburger. 2013. “The Progression Free Survival-Plateau with Vascular Endothelial Growth Factor Receptor Inhibitors – Is There More to Come?” European Journal of Cancer 49 (11): 2504–11.

Gujral, Taranjit S., Marina Chan, Leonid Peshkin, Peter K. Sorger, Marc W. Kirschner, and Gavin MacBeath. 2014. “A Noncanonical Frizzled2 Pathway Regulates Epithelial-Mesenchymal Transition and Metastasis.” Cell 159 (4): 844–56.

Gujral, Taranjit Singh, Leonid Peshkin, and Marc W. Kirschner. 2014. “Exploiting Polypharmacology for Drug Target Deconvolution.” Proceedings of the National Academy of Sciences of the United States of America 111 (13): 5048–53.

Harris, Leonard A., Peter L. Frick, Shawn P. Garbett, Keisha N. Hardeman, B. Bishal Paudel, Carlos F. Lopez, Vito Quaranta, and Darren R. Tyson. 2016. “An Unbiased Metric of Antiproliferative Drug Effect in Vitro.” Nature Methods 13 (6): 497–500.

Hsieh, James J., Mark P. Purdue, Sabina Signoretti, Charles Swanton, Laurence Albiges, Manuela Schmidinger, Daniel Y. Heng, James Larkin, and Vincenzo Ficarra. 2017. “Renal Cell Carcinoma.” Nature Reviews. Disease Primers 3 (March): 17009.

Jászai, József, and Mirko H. H. Schmidt. 2019. “Trends and Challenges in Tumor Anti-Angiogenic Therapies.” Cells 8 (9): 1102.

Kalinsky, Kevin, Fangxin Hong, Carolyn K. McCourt, Jasgit C. Sachdev, Edith P. Mitchell, James A. Zwiebel, L. Austin Doyle, et al. 2021. “Effect of Capivasertib in Patients with an AKT1 E17K-Mutated Tumor: NCI-MATCH Subprotocol EAY131-Y Nonrandomized Trial.” JAMA Oncology 7 (2): 271–78.

Khan, Kabir A., and Roy Bicknell. 2016. “Anti-Angiogenic Alternatives to VEGF Blockade.” Clinical & Experimental Metastasis 33 (2): 197–210.

Letai, Anthony. 2006. “Growth Factor Withdrawal and Apoptosis: The Middle Game.” Molecular Cell 21 (6): 728–30.

Liu, Minghui, Hongyu Liu, and Jun Chen. 2018. “Mechanisms of the CDK4/6 Inhibitor Palbociclib (PD 0332991) and Its Future Application in Cancer Treatment (Review).” Oncology Reports 39 (3): 901–11.

Luo, Ji, Michael J. Emanuele, Danan Li, Chad J. Creighton, Michael R. Schlabach, Thomas F. Westbrook, Kwok-Kin Wong, and Stephen J. Elledge. 2009. “A Genome-Wide RNAi Screen Identifies Multiple Synthetic Lethal Interactions with the Ras Oncogene.” Cell 137 (5): 835–48.

Mitchell, Scherise, Asuka Ota, William Foster, Bin Zhang, Zixing Fang, Shilpa Patel, Stanley F. Nelson, Steve Horvath, and Yibin Wang. 2006. “Distinct Gene Expression Profiles in Adult Mouse Heart Following Targeted MAP Kinase Activation.” Physiological Genomics 25 (1): 50–59.

Mollica, Veronica, Vincenzo Di Nunno, Lidia Gatto, Matteo Santoni, Marina Scarpelli, Alessia Cimadamore, Antonio Lopez-Beltran, et al. 2019. “Resistance to Systemic Agents in Renal Cell Carcinoma Predict and Overcome Genomic Strategies Adopted by Tumor.” Cancers 11 (6). https://doi.org/10.3390/cancers11060830.

Na, Yi Rang, Jung Won Kwon, Da Young Kim, Hyewon Chung, Juha Song, Daun Jung, Hailian Quan, et al. 2020. “Protein Kinase A Catalytic Subunit Is a Molecular Switch That Promotes the Pro-Tumoral Function of Macrophages.” Cell Reports 31 (6): 107643.

Nawabi, Homaira, Stephane Belin, Romain Cartoni, Philip R. Williams, Chen Wang, Alban Latremolière, Xuhua Wang, et al. 2015. “Doublecortin-Like Kinases Promote Neuronal Survival and Induce Growth Cone Reformation via Distinct Mechanisms.” Neuron 88 (4): 704–19.

Potente, Michael, Holger Gerhardt, and Peter Carmeliet. 2011. “Basic and Therapeutic Aspects of Angiogenesis.” Cell 146 (6): 873–87.

Rata, Scott, Jonathan Scott Gruver, Natalia Trikoz, Alexander Lukyanov, Janelle Vultaggio, Michele Ceribelli, Craig Thomas, Taran Singh Gujral, Marc W. Kirschner, and Leonid Peshkin. 2020. “An Optimal Set of Inhibitors for Reverse Engineering via Kinase Regularization.” Cold Spring Harbor Laboratory. https://doi.org/10.1101/2020.09.26.312348.

Sarkar, Siddik, and Mahitosh Mandal. 2009. “Growth Factor Receptors and Apoptosis Regulators: Signaling Pathways, Prognosis, Chemosensitivity and Treatment Outcomes of Breast Cancer.” Breast Cancer: Basic and Clinical Research 3 (August): 47–60.

Sherr, Charles J., David Beach, and Geoffrey I. Shapiro. 2016. “Targeting CDK4 and CDK6: From Discovery to Therapy.” Cancer Discovery 6 (4): 353–67.

Shiojima Ichiro, and Walsh Kenneth. 2002. “Role of Akt Signaling in Vascular Homeostasis and Angiogenesis.” Circulation Research 90 (12): 1243–50.

Sweeney, Christopher, Sergio Bracarda, Cora N. Sternberg, Kim N. Chi, David Olmos, Shahneen Sandhu, Christophe Massard, et al. 2021. “Ipatasertib plus Abiraterone and Prednisolone in Metastatic Castration-Resistant Prostate Cancer (IPATential150): A Multicentre, Randomised, Double-Blind, Phase 3 Trial.” Lancet 398 (10295): 131–42.

Veríssimo, André, Eunice Carrasquinha, Marta B. Lopes, Arlindo L. Oliveira, Marie-France Sagot, and Susana Vinga. 2018. “Sparse Network-Based Regularization for the Analysis of Patientomics High-Dimensional Survival Data.” Cold Spring Harbor Laboratory. https://doi.org/10.1101/403402.

Wang, Huawei, John Lapek, Ken Fujimura, Jan Strnadel, Bei Liu, David J. Gonzalez, Wei Zhang, et al. 2018. “Pseudopodium-Enriched Atypical Kinase 1 Mediates Angiogenesis by Modulating GATA2-Dependent VEGFR2 Transcription.” Cell Discovery 4 (1): 26.

Yang, Jie, Qing-Dong Shi, Tian-Bao Song, Gai-Feng Feng, Wei-Jing Zang, Chang-Hong Zong, and Ling Chang. 2013. “Vasoactive Intestinal Peptide Increases VEGF Expression to Promote Proliferation of Brain Vascular Endothelial Cells via the CAMP/PKA Pathway after Ischemic Insult in Vitro.” Peptides 42 (April): 105–11.

Zetterberg, A., and O. Larsson. 1985. “Kinetic Analysis of Regulatory Events in G1 Leading to Proliferation or Quiescence of Swiss 3T3 Cells.” Proceedings of the National Academy of Sciences of the United States of America 82 (16): 5365–69.

Zhao, Xiaocheng, Pavel Nedvetsky, Fabio Stanchi, Anne-Clemence Vion, Oliver Popp, Kerstin Zühlke, Gunnar Dittmar, Enno Klussmann, and Holger Gerhardt. 2019. “Endothelial PKA Activity Regulates Angiogenesis by Limiting Autophagy through Phosphorylation of ATG16L1.” ELife 8 (October). https://doi.org/10.7554/eLife.46380.

Zhou, L., X-D Liu, M. Sun, X. Zhang, P. German, S. Bai, Z. Ding, et al. 2016. “Targeting MET and AXL Overcomes Resistance to Sunitinib Therapy in Renal Cell Carcinoma.” Oncogene 35 (21): 2687–97.

